# Autocrine regulation of adult neurogenesis by the endocannabinoid 2-arachidonoylglycerol (2-AG)

**DOI:** 10.1101/2021.02.11.430731

**Authors:** Lena-Louise Schuele, Britta Schürmann, Andras Bilkei-Gorzo, Andreas Zimmer, Este Leidmaa

## Abstract

The endocannabinoid system modulates adult hippocampal neurogenesis by promoting the proliferation and survival of neural stem and progenitor cells (NSPCs). Specifically, deleting cannabinoid receptors on NSPCs or the constitutive deletion of the endocannabinoid 2-arachidonoylglycerol (2-AG) producing enzyme diacylglycerol lipase alpha (DAGLa) disrupts adult neurogenesis. However, it is not known which cells are the producers of 2-AG relevant to neurogenesis. In this paper, we investigated the cellular source of endocannabinoids in the subgranular zone (SGZ) of the hippocampus, an important neurogenic niche. For this purpose, we used two complementary Cre-deleter mouse strains to delete DAGLa either in neurons or astroglia and NSPCs. Surprisingly, neurogenesis was not altered in mice with a deletion of *Dagla* in neurons (Syn-Dagla KO), although they are the main source for the endocannabinoids in the brain. In contrast, mice with a specific inducible deletion of *Dagla* in NPSCs and astrocytes (GLAST-CreERT2-Dagla KO) showed a strongly impaired neurogenesis with significantly reduced proliferation and survival of newborn cells. These results identify *Dagla* in NSPCs in the SGZ of dentate gyrus or in astrocytes, as the cellular source for 2-AG in adult hippocampal neurogenesis. In summary, 2-AG produced by progenitor cells or astrocytes in the SGZ regulates adult hippocampal neurogenesis.

**Summary:** DAGLa in neuronal progenitor cells in the SGZ of dentate gyrus is identified as the cellular source for 2-AG in adult hippocampal neurogenesis.

## Introduction

Adult neurogenesis is a process in which new neurons are continuously generated in the mammalian adult brain (Altman and Das, 1965). In particular, two major regions containing neural stem and progenitor cells (NSPCs) were identified as neurogenic niches, the subgranular zone (SGZ) of hippocampal dentate gyrus (DG) and the subventricular zone (SVZ). Even though the biological role of adult neurogenesis is far from being fully elucidated, it is now well established that new adult-born neurons from SVZ and SGZ are functionally incorporated in the olfactory bulb or hippocampus, respectively, participating in the function of this areas (Bond et al., 2015).

Adult neurogenesis is regulated by a variety of different intrinsic and extrinsic factors. Several studies revealed that endocannabinoid signaling plays a crucial role in the regulation of proliferation, differentiation and survival of progenitor cells in the neurogenic niches. Specifically, cannabinoid receptor 1 (CB1) KO mice, lacking the main endogenous cannabinoid receptor of the CNS, were found to show highly decreased adult neurogenesis (Aguado et al., 2005). Similarly, diacylglycerol lipase alpha (*Dagla*) deficient mice, lacking the main producing enzyme of the most abundant cannabinoid 2-arachidonoylglycerol (2-AG) displayed impairments in adult neurogenesis (Gao et al., 2010; Jenniches et al., 2016). However, the cell type that regulates adult neurogenesis by endocannabinoids is not yet identified. Regarding CB1 signaling, a very recent study found that deleting CB1 specifically in NSPCs using an inducible Nestin-CreERT2 strain was sufficient to lead to reduced proliferation of progenitors in the DG of adult mice. The reduction of adult neurogenesis in this mouse line led to a concomitant depressive-like behavior of the mice, as accessed in the forced swim test (Zimmermann et al., 2018). Nevertheless, it is unknown where the endocannabinoids that activate CB1 receptors on progenitor cells come from. Elucidating this was the aim of this study. Therefore, conditional *Dagla* KO mice with a deletion in either neurons or astrocytes and NSPCs were used.

## Results

### Neural stem and progenitor cells (NPSCs) in subgranular zone (SGZ) of hippocampus express *Dagla*

Using a specific and sensitive *in situ* hybridisation method (RNAscope), we identified neural stem and progenitor cells (NSPCs) in the subgranular zone (SGZ) of the hippocampal dentate gyrus (DG) by the expression of *Sox2*. These cells also express *Dagla* (Figure 1), although the expression level seems to be somewhat lower compared to the cells in the granular layer. To validate this result on the protein level, we also performed immunostainings with a DAGLa specific antibody. NSPCs were identified by their localization, morphology and GFAP expression. As shown in Figure 3B, GFAP-positive cells in the SGZ of control mice were also positive for DAGLa. Together, these results demonstrate that NSPCs express DAGLa.

**Figure 1:**
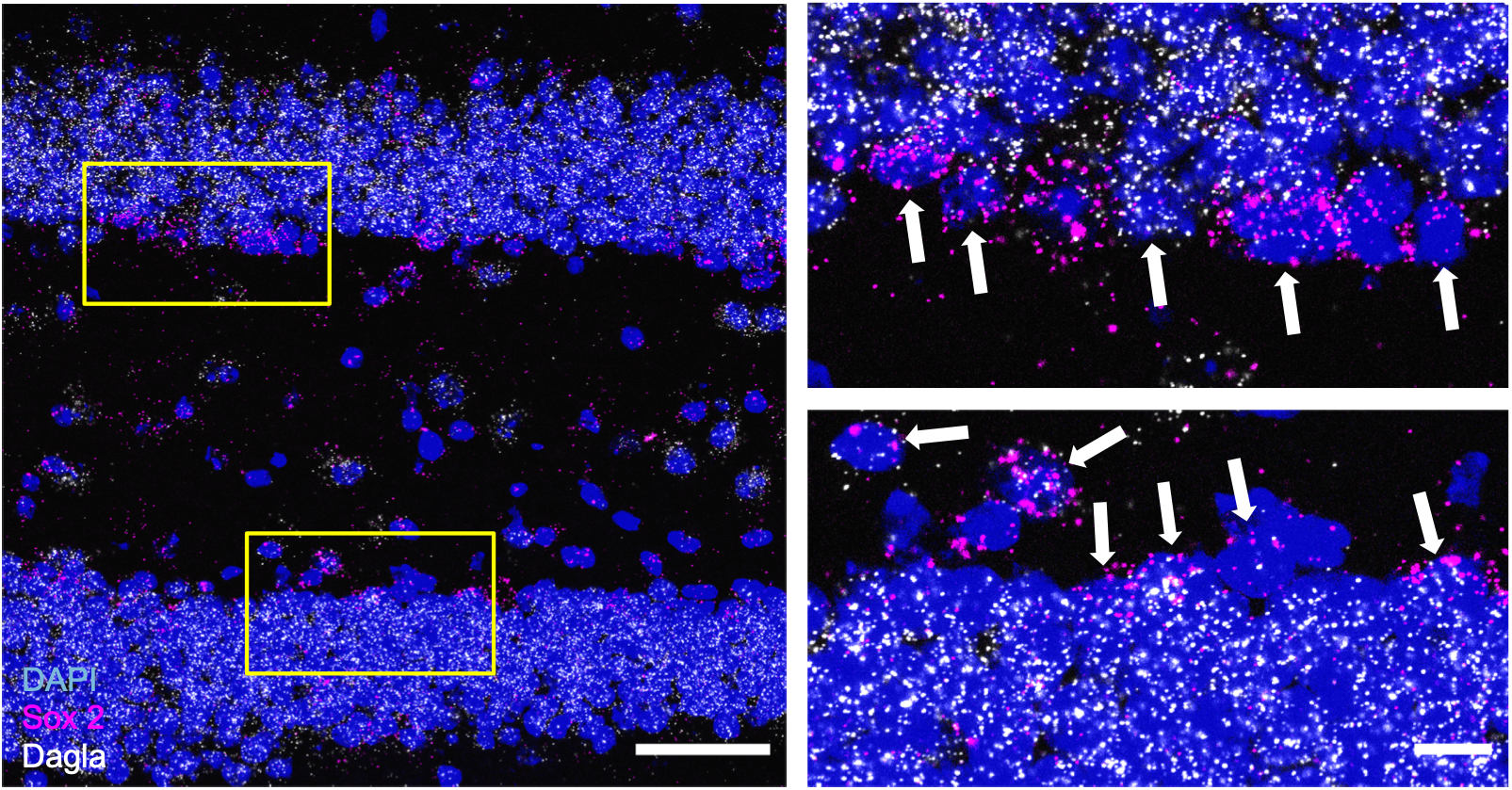
**(A)** Representative image of an RNAscope *in situ* hybridisation assay detecting transcripts of diacylglycerol lipase alpha *Dagla* (white) and neural progenitor marker *Sox 2* (magenta) in the dentate gyrus of hippocampus (DG) (scale bar 50 μm) **(B)** The magnified insets (scale bar 10 μm) show the subgranular zone (SGZ); the progenitor layer in the DG. White arrows indicate Sox 2-positive cells that co-express *Dagla* (scale bar 10 μm).

### Hippocampal deletion of DAGLa

The two conditional Cre-deleter strains used in this study (Syn-Cre and GLAST-CreERT2) showed very different but complementary Cre expression patterns in the hippocampal DG and *cornu ammonis* 3 (CA3) regions. Thus, as shown in Figure 2A using RosaTomato reporter mice, Syn-Cre mice specifically expressed Cre in virtually all neurons of the DG and CA3. In contrast, GLAST-CreERT2 mice showed no expression of tdTomato in neurons, but in astroglial cells (for quantification see Schuele et al., 2020). In the DG, cells expressing tdTomato were mainly NSPCs, because they were located in the neurogenic niche of the SGZ and showed the typical shape and GFAP expression of NSPCs (Figure 2B left). Using a co-staining with the proliferation marker Ki67, we found that 68% of the proliferating cells in the DG expressed tdTomato (Figure 2B right). This indicates that GLAST-CreERT2 is also active in NSCs and NPCs, confirming the findings previously reported by Mori et al., (2006). We therefore crossed Syn-Cre and GLAST-CreERT2 deleter strains with floxed *Dagla* (Dagla fl/fl) mice, thus deleting DAGLa in the corresponding cell populations. This was confirmed by immunohistochemisty (Figure 2C-D), which showed a moderate reduction of the DAGLa signal in tamoxifen-induced GLAST-CreERT2-Dagla KO mice. In Syn-Dagla KO mice the immunostaining of DAGLa was markedly reduced in the CA1 region and virtually abolished in the DG and CA3 areas, consistent with a deletion of DAGLa in DG neurons. The quantification of DAGLa signal intensity in the DG showed a reduction of 76.4% for Syn-Dagla KO and 23.9% for GLAST-Dagla KO. These observations were further validated using RNAscope *in situ* hybridisation. *Dagla* expression was still present in the SGZ in Syn-Dagla mice despite the deletion in all other DG cells. An opposing picture was observed in GLAST-Dagla mice, where *Dagla* was reduced in the progenitor cell layer but not in the rest of the DG (Figure 3A). A co-immunostaining with a marker for neurons (NeuN), marker for astrocytes and NPCs (GFAP) showed that the same holds true on the protein level. DAGLa is expressed in GFAP-positive NPCs in Syn-Dagla mice but not present in the neurons in granular cell layer, whereas the opposite is the case for GLAST-CreERT2-Dagla mice (Figure 3B). Thus, in Syn-Dagla mice *Dagla* was deleted from all DG cells, with the exception of NSPCs and few astrocytes, whereas, conversely, in GLAST-CreERT2-Dagla mice *Dagla* was deleted only in NSPCs and a few astrocytes.

**Figure 2:**
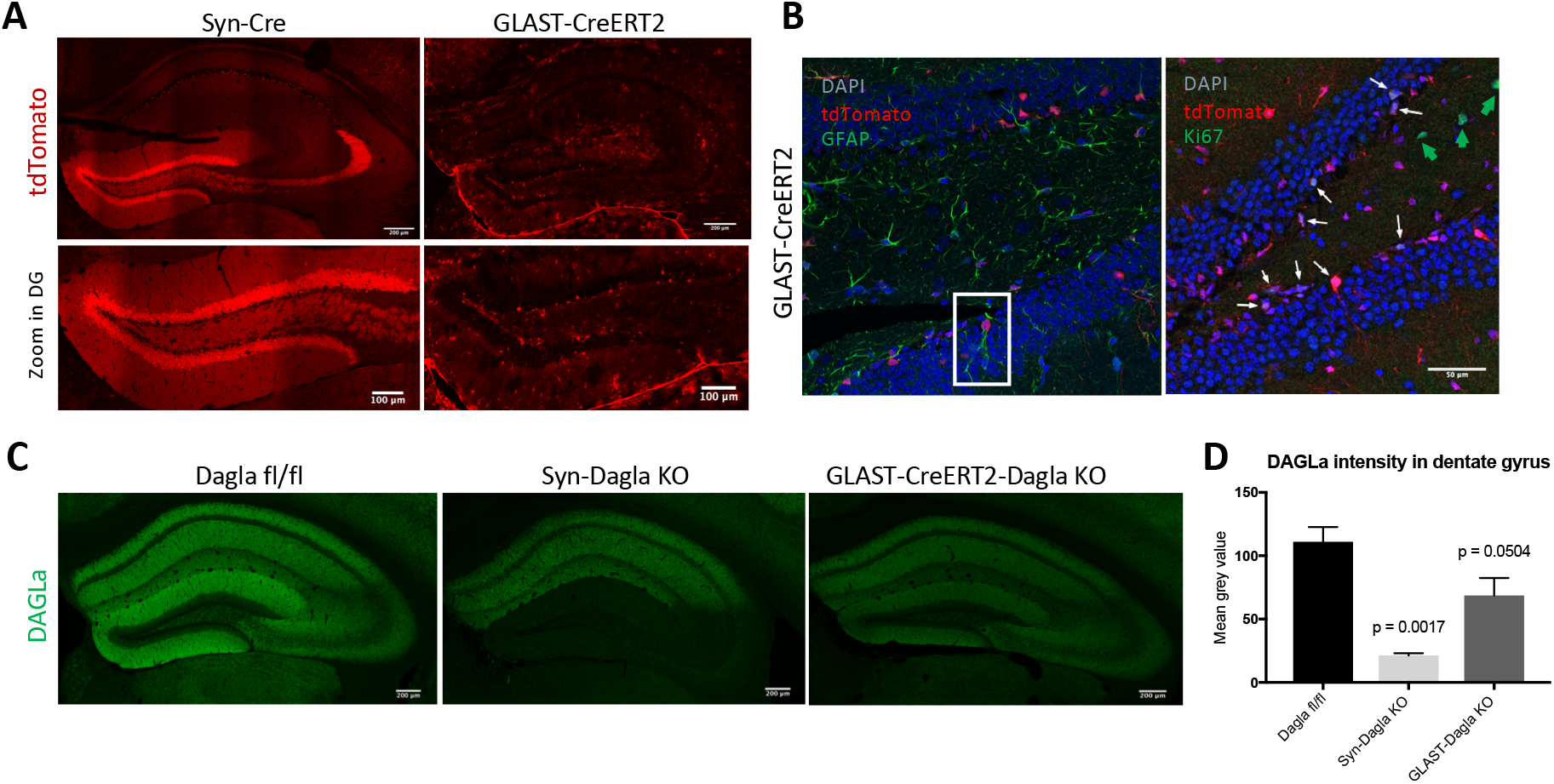
**(A)** Validation of Cre-efficacy in hippocampus (top, scale bar 200 μm) and dentate gyrus (DG) (bottom, scale bar: 100 μm) in Syn-Cre and tamoxifen-induced GLAST-CreERT2 lines crossed with the RosaTomato reporter line. GLAST-CreERT2 show tdTomato expression (red) and thus recombination activity in areas populated mostly by glial cells, especially in the subgranular zone (SGZ) of DG, but not in neurons in the granule cell layer of DG. Syn-Cre mice show tdTomato expression specifically in neurons in the granule cell layer of DG and CA3 but not in CA1 and CA2 or SGZ. **(B) (Right)** Co-staining of GLAST-CreERT2-RosaTomato with glial marker GFAP (green) in dentate gyrus. TdTomato expressing glia cells are mainly localized in the subgranular zone (SGZ) of dentate gyrus and show typical shape of neural stem and progenitor cells (NSPCs, example in white box)). Only few astrocytes not located in SGZ show tdTomato expression. **(Left)** Co-staining of GLAST-CreERT2-RosaTomato with proliferation marker Ki67 (green). 68% of all proliferating cells express Cre; 17% of all cre-expressing cells are proliferating cells (n=3 animals/group; white arrows: cells co-expressing tdTomato and Ki67; green arrow: proliferating cells not expressing tdTomato). **(C)** Representative images of DAGLa immunostainings from hippocampi. In Dagla fl/fl control mice, DAGLa is highly expressed throughout the hippocampus. Syn-Dagla KO mice show a very low DAGLa signal in dentate gyrus (DG) and cornu ammonis 3 (CA3), while CA1 and CA2 show similar DAGLa expression as in Dagla fl/fl control mice. The DAGLa signal in hippocampus of GLAST-CreERT2-Dagla KO mice is markedly reduced compared to control mice. **(D)** Quantification of DAGLa signal intensity in the DG of conditional knockout mice. GLAST-CreERT2-Dagla KO mice show a significant reduction of DAGLa signal in the dentate gyrus, that was even more pronounced in Syn-Dagla KO mice (1-way ANOVA: F_2,6_=18.34; p=0.0028; n=3 per group).

**Figure 3.**
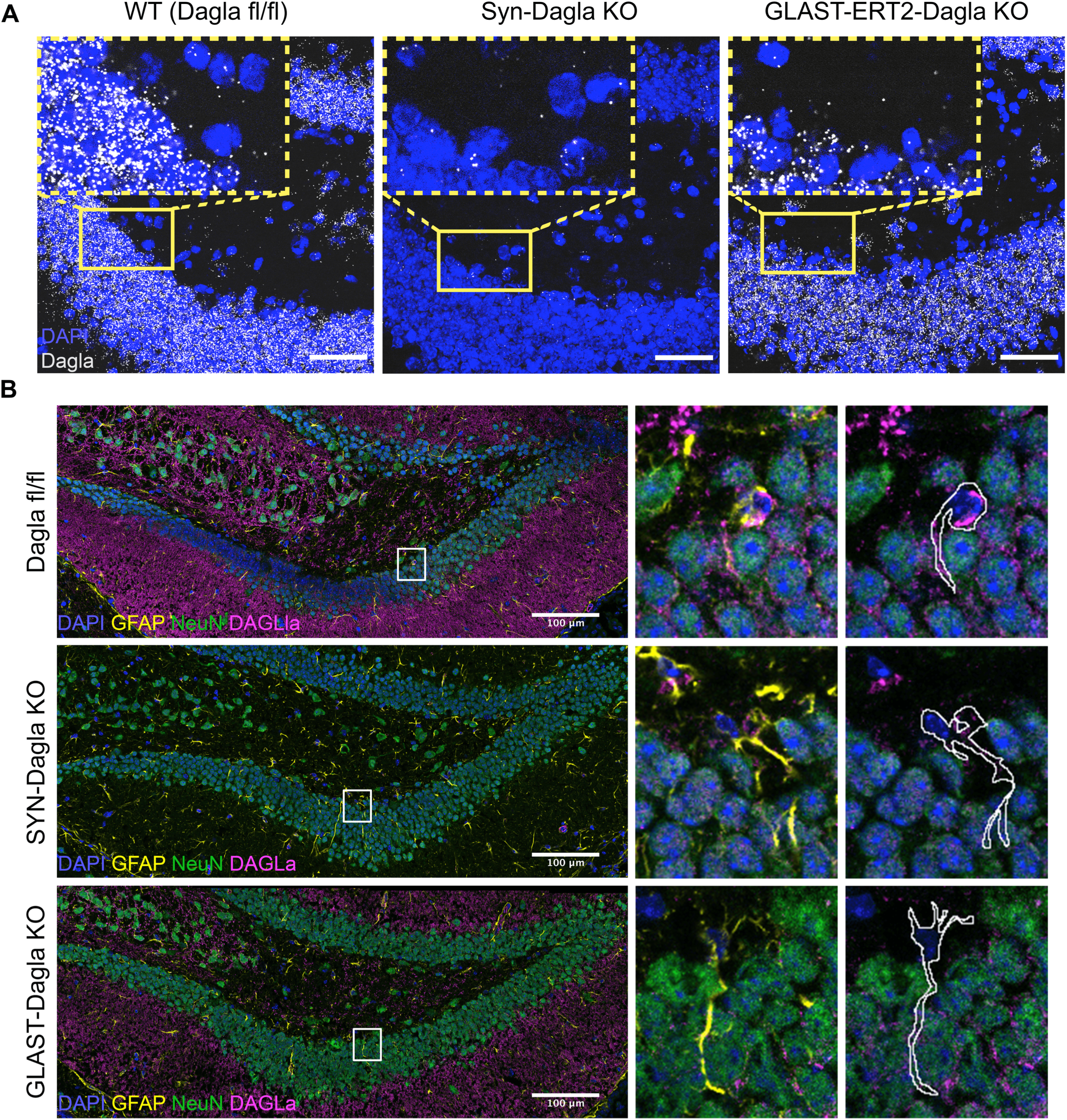
**(A)** Representative images of an RNAscope *in situ* hybridisation assay detecting transcripts of *Dagla* (white) in the DG in the hippocampus of Dagla fl/fl control, Syn-Dagla KO and GLAST-CreERT2-Dagla KO mice. *Dagla* is deleted in the neuronal granular cell layer in Syn-Dagla and in the progenitor-containing subgranular zone (SGZ) in GLAST-CreERT2-Dagla KO mice (scale bar 50 μm). The magnified insets (scale bar 10 μm) show SGZ with white arrows indicating *Dagla* expressing cells in Syn-Dagla KO and white arrowheads Dagla-negative cells in GLAST-CreERT2-Dagla KO mice in the NSC layer (n= 3-4 animals per group). **(B)** Immunostaining detecting DAGLa (magenta) co-localization with GFAP (yellow) and NeuN (green). The magnified insets show examples of GFAP-positive cells with NSPC morphology delineated with white. Progenitors express DAGLa in Dagla fl/fl control and Syn-Dagla KO SGZ and but not in GLAST-CreERT2-Dagla KO. NeuN positive neurons in granule cell layer express DAGLa in control and GLAST-CreERT2-Dagla KO sections, but not in Syn-Dagla KO (scale bar 10 μm).

### Adult hippocampal neurogenesis in conditional Dagla KO mice

GLAST-Dagla KO mice showed a strongly reduced number of BrdU positive cells in the DG one day after the last BrdU injection, suggesting decreased proliferation of NSPCs in this mouse line (Figure 4A-B). Also 21 days after the last BrdU injection that is used to estimate the survival of newly born cells, GLAST-Dagla KO mice showed less BrdU positive cells compared to littermate control mice. Please note that the ratio of BrdU positive cells on days 1 and 21 was similar in both genotypes, suggesting that the survival of NSPCs was not affected per se. Differentiation, measured by co-localization of BrdU with either a neuronal marker (NeuN) or astrocytic markers (GFAP or S100ß) was also unchanged (Figure 4C). Surprisingly, Syn-Dagla KO mice, with a complete deletion of *Dagla* in all DG neurons, showed neither changes in the proliferation nor in the survival of NSPC in the DG. As shown in Figure 5, the number of BrdU positive cells was similar to that observed in littermate control mice on day 1 and day 21 after BrdU injections. Analysis of co-localization of BrdU with cell specific markers also showed that the differentiation of neural progenitors was not changed in Syn-Dagla KO mice (Figure 5C).

**Figure 4.**
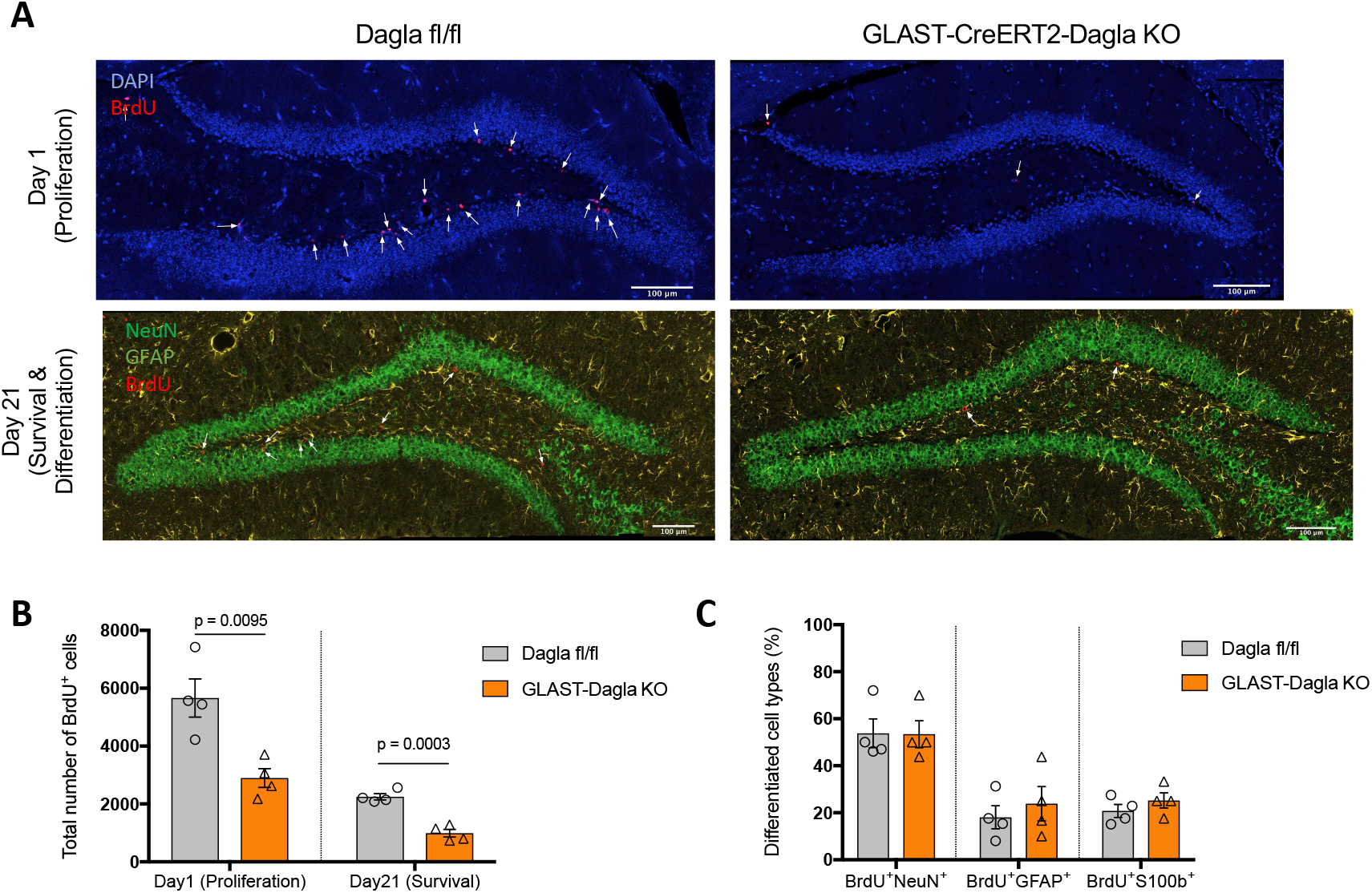
Adult hippocampal neurogenesis in GLAST-CreERT2-Dagla mice. **(A)** Representative immunohistochemistry micrographs of GLAST-CreERT2-Dagla WT and KO mice one or 21 days after BrdU injections in dentate gyrus to study proliferation and survival/differentiation, respectively (blue: DAPI; red: BrdU; green: NeuN; yellow: GFAP; scale bar: 100 μm). **(B)** The number of BrdU-positive cells in dentate gyrus of GLAST-CreERT2-Dagla KO mice was significantly lower compared to WT controls one day (p=0.0095) as well as 21 days (p=0.0003) after BrdU injections. **(C)** To analyze differentiation of progenitor cells, co-expression of BrdU-positive cells with neuronal marker (NeuN) or astrocytic marker (GFAP and S100) on day 21 were quantified. There were no changes in differentiation between GLAST-CreERT2-Dagla KO and WT control mice. Values represent mean ± SEM; n = 4 animals/group; 6 analyzed pictures per animal. Student’s t-test.

**Figure 5.**
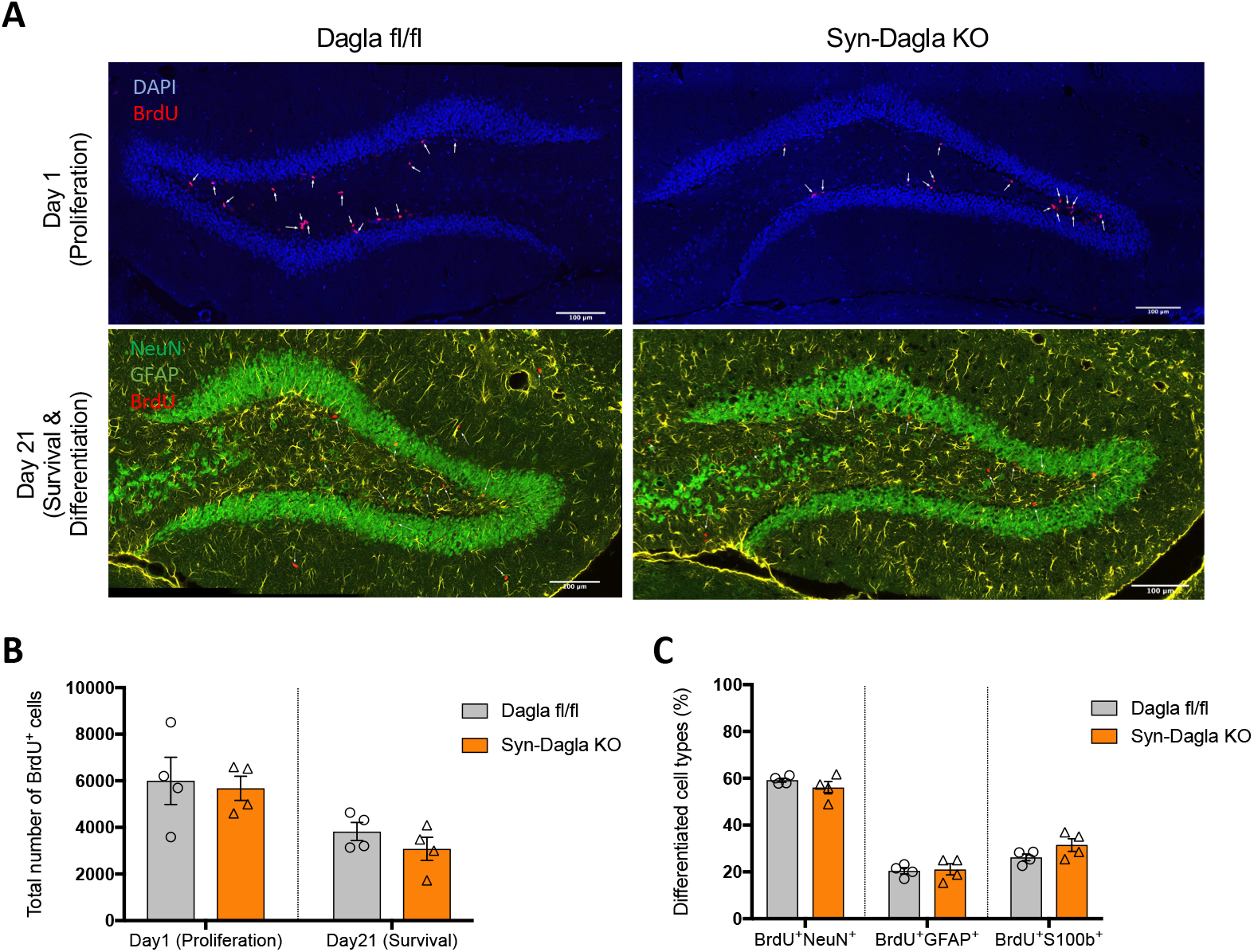
Adult hippocampal neurogenesis in Syn-Dagla mice. **(A)** Representative immunohistochemistry stainings of Syn-Dagla KO and control mice. Syn-Dagla KO mice show similar number of BrdU-positive cells (red) 1 day after the last injection. 21 days after BrdU injections, the number of BrdU positive cells does not differ between Syn-Dagla KO mice and controls. Additionally, brains were stained with an astrocytic marker GFAP (yellow) and a neuronal marker NeuN (green) to investigate differentiation, which was unchanged (scale bar: 100 μm). **(B)** The number of BrdU-positive cells in dentate gyrus of Syn-Dagla KO mice was similar to controls, one and 21 days after BrdU injections. **(C)** Differentiation of progenitor cells in dentate gyrus of Syn-Dagla KO mice. BrdU-positive cells were analyzed for co-expression of neuronal marker NeuN and astrocytic markers GFAP and S100beta. Values represent mean ± SEM; n = 4 animals/group, 6 analyzed pictures/animal. Student’s t-test.

## Discussion

In this study we show that the endocannabinoid 2-AG producing enzyme DAGLa is expressed in neural stem and progenitor cells (NSPCs) of the hippocampal dentate gyrus (DG) region, albeit at lower levels compared to neurons in the granule cell layer. Nevertheless, DAGLa expression in NSPCs is important for the regulation of adult hippocampal neurogenesis, whereas 2-AG produced by neurons in the DG surprisingly does not affect neurogenesis. Thus, deleting *Dagla* from NSPCs in the DG using inducible GLAST-CreERT2-Dagla KO mice, resulted in a prominent deficit in the proliferation of newborn cells. In contrast, neurogenesis was completely normal when *Dagla* was deleted from DG neurons using Syn-Dagla KO mice.

We have recently quantified *Dagla* expression in many brain regions in wild type mice, as well as in the GLAST-CreERT2-Dagla KO strain used here (Schuele et al., 2020). We observed a specific deletion of *Dagla* in the GLAST-CreERT2-Dagla KO strain in astrocytes but not in neurons, in almost all brain regions. An exception was the hilus of dentate gyrus (DG) where no astrocytic *Dagla* deletion was found in this mouse line. Our further analysis in the present paper demonstrated that *Dagla* was nevertheless deleted in the progenitor cell layer (NSPCs) of the DG, but not in other DG cells in GLAST-CreERT2-Dagla KO mice. The second mouse line used in the current study, Syn-Dagla KO mice, showed a complementary pattern: a complete deletion of *Dagla* from all neurons in the DG, but an intact *Dagla* expression in NSPCs. These fortuitous opposing patterns enabled us to precisely interrogate the cellular source of 2-AG in the regulation of adult hippocampal neurogenesis.

The fact that hippocampal neurogenesis was completely normal in Syn-Dagla KO mice was striking and contrary to our expectations. It is well-known that adult neurogenesis is influenced by physical activity, environmental factors and emotional states. Thus, environmental enrichment or running wheel activity can stimulate the proliferation and/or survival of NSPCs, whereas stress and ageing reduce progenitor cell proliferation (Dubreucq et al., 2010; Baptista and Andrade, 2018; Shohayeb et al., 2018). Many studies have demonstrated that these factors modulate not only adult neurogenesis, but also affect cognitive functions, providing a link between adult neurogenesis and cognition and behavior. The mechanisms of how these environmental stimuli affect neurogenesis are not yet completely understood (Shohayeb et al., 2018), but seem to involve the neuronal activity-driven release of growth factors such as BDNF and FGF2. Considering the prominent role of the endocannabinoid system in the modulation of neuronal activity, we assumed that endocannabinoids would indirectly modulate NSPC proliferation and survival through their production by neurons as a result of neuronal activity (Freund et al., 2003; Kano et al., 2009). Contrary to this hypothesis, our present results strongly suggest, however, that endocannabinoids modulating NSPC proliferation originate from progenitors themselves. Our findings are entirely consistent with a recent report showing that cannabinoid receptor 1 (CB1) in NSPCs is also important for NSPC proliferation (Zimmermann et al., 2018). In this study CB1 was specifically deleted in NSPCs using a Nestin-CreERT2 deleter strain, whereas CB1 expression in DG neurons remained intact. These mice also showed a decreased proliferation of NSPCs (Zimmermann et al., 2018). Therefore, deleting the CB1 receptor in NSPCs, or the main ligand-producing enzyme DAGLa in NSPCs, produced a very similar phenotype. Together, these findings suggest that an endocannabinoid-mediated autocrine signaling mechanism in NSPCs modulates adult hippocampal neurogenesis.

A similar autocrine mechanism has also been suggested during the development of the nervous system. DAGLa and CB1 are both expressed in migratory neuroblasts throughout the cell, including the tip of the filopodia in the terminal growth cone-like structure (Oudin et al., 2011). There is accumulating evidence to suggest that this autocrine mechanism contributes to the regulation of axonal growth and guidance, probably by acting downstream of a cell adhesion molecules/fibroblast growth factor receptor (CAM/FGFR) signaling system (Oudin et al., 2011). Specifically, FGFR activation induces 2-AG production by sequential activation of phospholipase Cγ (PLCγ) which produces diacylglycerol, and then 2-AG through DAGLa. This in turn results in CB1 activation in neuroblasts and modulates cortical neuron specification, elongation and morphological differentiation during embryonal development (Williams et al., 2003; Galve-Roperh et al., 2013; Maccarrone et al., 2014).

Autocrine manner of cannabinoid receptor activation by 2-AG also has been suggested to happen in the adult brain (Oudin et al., 2011). Oligodendrocyte precursors also produce 2-AG and express CB1 and CB2 receptors and basal tone of 2-AG and cannabinoid receptors maintains the precursor proliferation in culture (Gomez et al, 2015), again hinting towards a possible autocrine mechanism regulating proliferation.

It is currently not known which signaling mechanisms act upstream of the endocannabinoid system in the modulation of adult neurogenesis. In addition to growth factor receptor activation these may include metabotropic receptors, such as dopamine receptor D1 which has been shown to regulate DAGLa activity (Shonesy et al., 2020) as well as NSPC proliferation (Takamura et al., 2014) or the well-established upstream regulator of DAGLa, the glutamate receptor. We propose that an autocrine endocannabinoid signaling in NSPCs may either serve in signal potentiation or, alternatively, function as a signal integrator, for example for coincidence detection of several neurogenic signals. In this context, it is interesting to note that endocannabinoids have biphasic effects on self-renewal in other types of stem cells. Thus, low doses stimulate proliferation of immune stem cells and high doses inhibit it (Guzman et al, 2002).

The field of stem cell biology has gained attention and made progress in the recent decades because it holds great promise to repair and rejuvenate different tissues in the adult organism. Endocannabinoid system is a possible target for understanding these processes as it regulates stem cell biology in the whole organism throughout life-span (Galve-Roperh et al, 2013). In early embryogenesis, endocannabinoids influence embryonic and trophoblast stem cell differentiation and survival. Its effects continue in ectoderm-derived neural tissues and mesoderm-derived hematopoietic and mesenchymal stem cells, thus affecting blood cells, adipose tissue, bone, muscle and epithelia (Galve-Roperh et al, 2013; Maccarone et al. 2014). Considering this, it is not surprising that total *Dagla* or CB1 knockouts show reduced breeding efficiency (smaller litters, less offspring). It is remarkable, however, that mice lacking *Dagla* or *Cb1* do not have gross developmental deficits (Jenniches et al., 2016). A plausible explanation is that there is a redundancy of pathways and compensation by other factors regulating these crucial developmental stages. In the adulthood, however, there seems to be less plasticity and potential for compensation. As exemplified by the fact that deletion of *Dagla* in a subpopulation of neural progenitors in our study leads to an almost 50% decrease of proliferation. This becomes important in natural aging processes in which 2-AG levels decrease (Piyanova et al., 2015) and there is a depletion of NSPC in the hippocampus possibly leading to cognitive deficits. In fact, the restoration of cognitive function in old mice by chronic low dose of Δ9-tetrahydrocannabinol (Bilkei-Gorzo et al., 2017) may depend at least partly on its effects on neurogenesis. Moreover, the same treatment could be also used to ameliorate depression in elderly people.

We previously demonstrated that GLAST-CreERT2-Dagla KO mice had a profound impairment of maternal behavior and a depressive-like behavioral phenotype (Schuele et al., 2020). It is well known that interfering with adult hippocampal neurogenesis can results in depressive-like behavioral changes (Anacker and Hen, 2017). Conversely, many animal models of depression show impaired hippocampal neurogenesis (Du Preez et al., 2021). It is thus possible that the behavioral phenotype of GLAST-CreERT2-Dagla KO mice was at least partially due the impaired hippocampal neurogenesis. This idea is supported by a study of Zimmermann et al. (2018), which showed that mice lacking CB1 on NSPCs also have a depression-like phenotype and reduced adult hippocampal neurogenesis (Zimmermann et al. 2018).

Together, we propose that progenitor cells produce 2-AG which binds to their own CB1 receptors and promotes adult neurogenesis. Impairments in this process might eventually lead to development of depressive-like behaviors.

## Methods

### Animals

Dagla fl/fl (B6.cg(Dagla)tm1Zim) mice (Jenniches et al., 2016) mice on a C57BL/6J genetic background (2-5 month old) were used in this study. These mice were crossed to mouse lines expressing Cre recombinase either under the control of a neuron specific synapsin (Syn) promoter, or under the control of an astrocyte-specific glutamate transporter promoter (GLAST). The synapsin-Cre line was obtained from The Jackson Laboratory B6.Cg-Tg(Syn1cre)671Jxm/J). The glutamate transporter-Cre line (B6.Cg-Slc1a3tm1(cre/ERT2)Mgoe) was generated by Mori et al., 2006 (Mori et al., 2006). This strain has an insertion of a tamoxifen-inducible form of Cre (CreERT2) in the locus of astrocyte-specific glutamate aspartate transporter (GLAST, Slc1a3tm1). The resulting mouse lines are referred to as Syn-Dagla and GLAST-CreERT2-Dagla, respectively. Mice were always bred heterozygously for Cre. Littermates not expressing the Cre gene, designated as “Dagla fl/fl”, were used as controls. The floxed *Dagla* allele, the wild type allele, the knocked out *Dagla* allele, and the Cre locus were identified by polymerase chain reaction (PCR) using appropriate primers (Cre1_fwd CATTT GGGCC AGCTA AACAT, Cre2_fwd GCATT TCTGG GGATT GCTTA, Cre1_rev CCCGG CAAAA CAGGT AGTTA, Cre2_rev TGCAT GATCT CCGGT ATTGA, Dagla_fwd TAG CTT AGC CCC CAT GTG AC, Dalga WT rev GAG ATG GGT TCC ACC TCC TT, Dagla fl/KO_rev CCC AGT AGC CAC AGA ACC AT or CGC AGC CCA AAA GAT ACA AT). To validate region specific Cre expression we used RosaTomato reporter mice (The Jackson Laboratory (B6.Cg-Gt(ROSA)26Sortm14(CAG-tdTomato)Hze/J). They carry a reporter construct in the ubiquitously expressed Gt(ROSA)26Sor locus. The reporter is expressed after Cre-mediated excision of a loxP-flanked STOP cassette. We used mice that were heterozygous for RosaTomato (wt/ins) and Cre (wt/tg). Mice were housed under a 12 hours light/dark cycle (lights on 9am-9pm) with ad libitum access to food and water. All experiments were approved by the North Rhine-Westphalia State Environment Agency (AZ: 84-02.04.2017.A234).

### RNAscope assay

The RNAscope method was used to evaluate the expression of Dagla in the cells of interest (www.acdbio.com/RNAScope). For this purpose, mice (n = 2-4 per genotype) were killed by decapitation, brains were quickly removed, flash frozen in dry ice-cooled isopentane and stored at −80 °C. Brains were cryosectioned at a thickness of 10 μm and mounted on SuperFrost Plus slides (Thermo Fisher, Schwerte, Germany). The exact Bregma coordinates were identified from every twelfth slide according to Paxinos Brain atlas (Franklin and Paxinos, 2008). RNAscope Multiplex Fluorescent Reagent kit was used according to manufacturer’s instructions (Advanced Cell Diagnostics, Newark, CA). The probes for *Dagla* mRNA (cat.no 478821) were multiplexed with the probes detecting the neural stem cell marker *Sox2* (cat. no 402041-C3). Confocal images of sections were obtained using a Leica TSC SP8 with 40x magnification (291 μm2 area) and analyzed using Fiji software (version 2.0.0-rc-69; NIH, Bethesda, MD, USA). Micrographs from at least two consecutive sections from an animal were analyzed per brain region (Bregma −1.34- −1.46 mm). In every assay, sections from constitutive Dagla KO mice (Jenniches et al., 2016) were used as controls for non-specific binding of the Dagla probe.

### Immunohistochemistry (IHC)

Mice were anesthetized with isoflurane and transcardially perfused with 20 mL of cooled PBS, followed by 60 mL of 4% formaldehyde. The brains were postfixed in 4% formaldehyde (only for co-stainings with tdTomato) for one hour, incubated in 20% sucrose/PBS for approximately 3 days, and frozen in dry ice-cooled isopentane. Tissues were stored at −80°C until further processing and cut in a cryostat with a chamber temperature of −22 °C. Slices were collected on SuperFrost Plus adhesion slides (Thermo Fisher) and stored at −20 °C. For DAGLa staining, brains were cut in 40 μm thick free-floating sections (Bregma −0.94 to −3.34 mm). For immunostainings with cell type specific markers and tdTomato, brains were cut into 16 μm slices (Bregma −1.46 until −2.46, every 8th section collected on a slide). For DAGLa co-staining with cell-specific markers, fresh frozen brains were cut in 10 μm thick slices (Bregma −1.34- - 1.46) and post-fixed for 20 minutes in 4% formalin in phosphate buffer before staining. After immunostaining, brain slices were briefly washed in ddH_2_O and embedded in DAPI Fluoromount-G(R) media (Southern Biotechnology Associates. Inc., Birmingham, AL). When costaining of DAGLa with cell-specific markers was performed, a mounting media ProLong™ Diamond Antifade with DAPI (ThermoFisher) was used. Fluorescence images were obtained with the Leica SP8 confocal microscope, with 20x (NA 0.07, pixel size: 0.001 mm x 0.001 mm) or 40x (NA 1.1, pixel size: 0.142 μm x 0.142 μm) objective lenses. Phosphate buffer (PB, 0.2 M, pH 7.6) was used as a diluent and wash solution for DAGLa staining (DGLa-Rb-Af380; Frontier Institute Co., Ltd., Hokkaido, Japan). Triton (0.3 %) was used for permeabilization (10 min). Nonspecific antibody binding was blocked with 5% normal goat serum and 0.2% Triton (1 hour at RT), DAGLa antibodies were incubated overnight at RT (1:400 in blocking solution). Secondary goat anti-rabbit Alexa-Fluor647 (Life Technologies, Darmstadt, Germany) was incubated for 2 hours at RT (1:500 in blocking solution). For DAGLa co-stainings with neuronal or astrocyte markers guniea pig anti-NeuN (1:1000 266004, Synaptic Systems) and chicken anti-GFAP (1:1000 ab4674, Abcam, Cambridge, GB). As secondary antibodies, goat anti-chicken Alexa488, goat anti-guniea pig Alexa568 and goat anti-rabbit Alexa647 were used. For tdTomato IHC phosphate-buffered saline (PBS; 0.01 M, pH 7.4) served as a general diluent and wash solution. Sections were permeabilized by 0.5% TritonX100 in PBS for one hour. Unspecific binding was blocked by 3% BSA for two hours prior to primary antibody incubation (24 hours at 4°C). Microglia were stained with rabbit anti-Iba1 antibody (1:2000 in 0.3% BSA/PBS; 019-19741, Wako, Richmond, VA), astrocytes with rabbit-anti-S100ß (1:2000 ab41548, Abcam) and chicken anti-GFAP (1:2000 ab4674 Abcam), neurons with rabbit anti-NeuN (1:500 ABN78C3, Merck, Millipore, Burlington, USA). Secondary antibodies were anti-rabbit Alexa-Fluor 488 (1:1000), donkey anti-rabbit Alexa-Fluor 647 (1:2000) or goat anti-chicken Alexa-Fluor 488 (1:2000).

### BrdU injections and IHC

To label proliferating cells, two months old mice (three months old in case of GLAST-Dagla and controls (tamoxifen injection at 2 months of age) were injected intraperitoneally with 5-bromo-2’-deoxyuridine (BrdU; 50 mg/kg bodyweight, dissolved in sterile saline) once a day for three consecutive days. To investigate proliferation of progenitor cells, 4 mice of each group were killed 24 h after the last BrdU injection. To determine differentiation and survival of progenitor cells, animals were killed after 21 days. For this, they were anesthetized with isoflurane and transcardially perfused with cooled PBS, following 4% PFA. Brains were postfixed in 4% PFA for 24 hours at 4°C. Brains were cryoprotected in 20% sucrose for three days at 4°C before they were frozen in dry ice cooled isopentane and stored at −80°C. Subsequently the entire hippocampus (Bregma −0.94 to −3.34 mm) was cut into 48 serial coronal free-floating cryosections of 40 μM thickness. Slices were stored in 48 well plates filled with cryoprotectant (50% PBS, 25% glycerol, 25% ethylenglycol, 0.025% sodiumazide) at −20°C. For immunohistochemical staining, every eighth hippocampal slice of each mouse was selected and transferred in one well of a 24-well-plate containing Netwell™ inserts (Costar). All following steps of the staining were done at room temperature under shaking, if not specified otherwise. First, slices were washed three times in TBS for five min to remove remaining cryoprotectant, followed by incubation in permeabilization buffer (TBS containing 0.3% Tween) for 10 minutes. After another washing step, slices were incubated in 2xSSC for 20 minutes at 65°C for antigen retrieval and then shortly washed in ddH_2_O. Since BrdU is located in the DNA, slices were incubated in 2M HCl for 30 min at 37°C to make DNA accessible for the antibody. To neutralize pH, slices were incubated in borate buffer for 10 min before they were washed in TBS again. Subsequently, slices were blocked in TBS plus II (TBS containing 0.3% Triton X-100, 5% goat serum, 2% BSA) for 1 hour.

To investigate the proliferation of progenitor cells, brain slices were incubated with rat anti-BrdU antibody (1:500 ab6326, Abcam in TBS plus II) overnight at 4°C under shaking. To investigate the survival and differentiation of progenitor cells, brain slices of mice were also incubated with a mouse NeuN (1:500, conjugated with AF488 MAB377X, Merck) and a rabbit GFAP antibody (1:1000 ab7260, Abcam). Since GFAP is also expressed in young neurons, we also used a S100beta antibody (1:2000 ab41548, Abcam) instead of the GFAP antibody. In both cases, brain slices were incubated in the antibody solutions over night at 4°C under shaking.

On the next day, slices were washed and then blocked in TBS plus II for 1 hour. As secondary antibody, goat-anti-rat AF594 (1:500 in TBS plus II) was used to label BrdU antibodies and goat-anti-rabbit AF647 (1:1000 in TBS plus II) to label GFAP-or S100beta-antibodies. Slices were incubated in the secondary antibody solution for 2 hours at RT. After another washing step slices were transferred to coverslips and embedded in DAPI Fluoromount-G^®^ (Southern Biotechnology Associates, Inc.). Images of the dentate gyrus region were obtained with Leica SP8 Confocal microscope with 40x objective in tile-scan mode.

To estimate all BrdU-positive cells in the entire hippocampus, the following calculation was used:

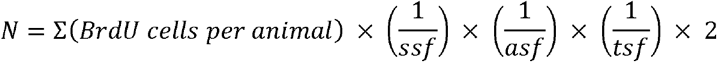

The selected sampling fraction (ssf) is 0.125 (6/48), because 6 out of 48 hippocampal slices were analyzed. Since the entire dentate gyrus was used as counting frame, the value for the area sampling fraction (asf) equals 1. Due to the thickness of slices (40 μm), the optical section of confocal microscope (1.3 μm) and an average diameter of nuclei of ~8 μm, the thickness of sampling fraction (tsf) was considered as 0.25 ((1.3 x 8)/40). Last, the calculated number of BrdU-positive cells was multiplied with two, because only one hemisphere was imaged for analysis.

Since the staining for survival and differentiation (D21) was always performed two times (with GFAP or with s100ß) on 6 slices, the average amount of BrdU-positive and NeuN-positive cells per mouse of both stainings was used for analysis. Differentiation was evaluated by determination of overlaps of BrdU signal and either neuronal- (NeuN) or astrocytic (GFAP/S100ß) markers.

## Author contributions

LLS, AZ, ABG, EL were involved in designing research studies. LLS, BS, ABG and EL conducted the experiments, acquired and the analyzed data. AZ provided the reagents. LLS, AZ and EL were writing the manuscript.

## Acknowledgements

This research leading to these results was funded by the Deutsche Forschungsgemeinschaft (DFG, German Research Foundation) - project number 324087152, and to AZ under Germany’s Excellence Strategy – EXC2151 – 390873048. We thank our lab members Joanna Komorowska-Müller, Alessandra Gargano, Eva Drews, Edda Erxlebe, Hanna Schrage, Kersin Nicolai, Anne Zimmer, and previous lab members Eva Beins, and Imke Jenniches for their help and constructive discussions.

## Notes

### Competing Interest Statement

The authors have declared no competing interest.

